# A Novel Explainable Fuzzy Clustering Approach for fMRI Dynamic Functional Network Connectivity Analysis

**DOI:** 10.1101/2023.01.29.526110

**Authors:** Charles A. Ellis, Robyn L. Miller, Vince D. Calhoun

**Affiliations:** Tri-institutional Center for Translational Research in Neuroimaging and Data Science: Georgia State University, Georgia Institute of Technology, Emory University, Atlanta, GA 30303 USA

## Abstract

Resting state functional magnetic resonance imaging (rs-fMRI) dynamic functional network connectivity (dFNC) analysis has illuminated brain network interactions across many neuropsychiatric disorders. A common analysis approach involves using hard clustering methods to identify transitory states of brain activity, and in response to this, other methods have been developed to quantify the importance of specific dFNC interactions to identified states. Some of these methods involve perturbing individual features and examining the number of samples that switch states. However, only a minority of samples switch states. As such, these methods actually identify the importance of dFNC features to the clustering of a subset of samples rather than the overall clustering. In this study, we present a novel approach that more capably identifies the importance of each feature to the overall clustering. Our approach uses fuzzy clustering to output probabilities of each sample belonging to states and then measures their Kullback-Leibler divergence after perturbation. We show the viability of our approach in the context of schizophrenia (SZ) default mode network analysis, identifying significant differences in state dynamics between individuals with SZ and healthy controls. We further compare our approach with an existing approach, showing that it captures the effects of perturbation upon most samples. We also find that interactions between the posterior cingulate cortex (PCC) and the anterior cingulate cortex and the PCC and precuneus are important across methods. We expect that our novel explainable clustering approach will enable further progress in rs-fMRI analysis and to other clustering applications.

## I. INTRODUCTION

Many resting state functional magnetic resonance imaging (rs-fMRI) studies have sought to gain insight into a variety of neuropsychiatric disorders and cognitive functions via the extraction and subsequent clustering of static and dynamic functional network connectivity (sFNC and dFNC) [1]–[3]. Several recent studies have presented methods to identify the key brain network interactions most responsible for the identified clusters [4]–[6]. However, this line of research has several key shortcomings. Namely, most existing studies use hard clustering methods that do not effectively account for within cluster variation. Second, this hard clustering has led to the use of explainability methods for FNC clustering that involve perturbation and require samples to completely switch clusters for a given feature to be considered important [4], [5] Given that only a minority of samples switch clusters following perturbation [4], [5], these methods give insight not into the features most important to the overall clustering but rather into the features to which a small minority of samples are highly sensitive. In this study, we present a novel FNC clustering approach for insight into neuropsychiatric disorders and accompanying explainability approach for insight into the brain network interactions important to the identified clusters.

In recent years, many rs-fMRI studies have sought to better understand neurological [1] and neuropsychiatric [2], [7] disorders and cognitive functions [6], [8] through the extraction and clustering of FNC data. Static FNC is a measure of the correlation between brain regions throughout a recording [4], [6], and dFNC is a measure of the correlation between brain regions in short windows over time [2]. These clustering analyses typically involve k-means clustering, a popular hard clustering method. In sFNC, subgroups of individuals are identified, and in dFNC, transitory states of neurological activity are identified. A subsequent analysis is then typically performed with the goal of understanding the relationship between the identified subgroup or state and the disorder or cognitive function of interest.

While cluster centroids are often visualized for insight into the features differentiating each identified subgroup or state [1]–[6], recent studies have sought to provide explainability methods capable of identifying the relative importance of each FNC feature to the clusters [4]–[6]. These methods build upon permutation feature importance [9], which was originally developed for supervised machine learning explainability, and involve perturbing a given feature and examining the percentage of samples that switch clusters. Relative to earlier efforts that used high numbers of statistical tests to identify differences in specific features on a univariate basis [10], these new explainability methods account for the multivariate nature of the original clustering when providing an importance estimate. Nevertheless, these methods do have a key weakness that arise from their use with hard clustering methods.

The use of hard clustering methods like k-means clustering involves a critical assumption. Hard clustering methods assign samples to only one cluster, regardless of whether samples bear a degree of similarity to other centroids. As such, samples very near to one another in a data space might be assigned to different centroids. This has also led to an issue in perturbation-based explainability approaches. Namely, only a small number of samples are so affected by perturbation that they actually switch clusters [4], [6]. This indicates that methods which quantify importance as the percent of samples that switch clusters following perturbation are only showing the sensitivity of a small minority of samples to perturbation, rather than the relative importance of a given feature to the overall clustering.

In this study, we present Global Permutation-based Distribution Divergence (GP2D), a novel approach that can be used to gain greater insight into the importance of features to the overall FNC clustering. Specifically, we use fuzzy c-means clustering, an approach that yields probabilities of each sample belonging to each cluster, and adapt existing explainability methods to calculate the divergence in the probability of samples belonging to each cluster following perturbation. We present our approach within the context of dFNC default mode network (DMN) analysis, identifying fuzzy states that uncover differences in activity between healthy controls (HCs) and individuals with schizophrenia (SZ). We then compare two variations of GP2D to Global Permutation Percent Change (G2PC) feature importance [4] and show that our approach captures the effects of perturbation across most samples, not only those that are sensitive enough to perturbation to switch clusters.

## II. METHODS

In this section, we describe our study approach. (1) We extract dFNC from a dataset composed of individuals with SZ and healthy controls. (2) We cluster the samples using fuzzy c-means. (3) We extract features related to the dynamics uncovered by the clustering. (4) We analyze the relationship between the extracted cluster-based features and SZ diagnosis. (5) We use an existing clustering explainability approach for insight into the dFNC features that are most important to the clustering. (6) We propose a novel clustering explainability method, and (7) we compare our explainability approach with the existing approach.

### A. Description of Data and Preprocessing

In this study, we used 151 rs-fMRI recordings from SZs and 160 from HCs that are part of the Functional Imaging Biomedical Informatics Research Network (FBIRN) dataset [11]. It has been used in a variety of neuroimaging studies [12], [13]. Data collection was performed at 7 sites: the University of Iowa, the University of New Mexico, the University of Minnesota, Duke University/the University of North Carolina at Chapel Hill, the University of California at San Francisco, the University of California at Irvine, and the University of California at Los Angeles. Institutional review boards at each site approved data collection procedures, and participants gave written informed consent.

The first 5 mock scans were removed before preprocessing. We used statistical parametric mapping (SPM12, https://www.fil.ion.ucl.ac.uk/spm/) and corrected for head motion via rigid body motion correction. We spatially normalized the data to an echo-planar imaging template in the standard MNI space and resampled to 3×3×3 mm^3^. We used a 6mm full width at half maximum Gaussian kernel to smooth the data. We then extracted independent components (ICs) using the Neuromark pipeline of the GIFT toolbox (http://trendscenter.org/software/gift). While the pipeline extracts 53 ICs from the gray matter of a number of brain networks, we only examined 7 ICs from the default mode network (DMN): 3 precuneus (PCN), 2 anterior cingulate cortex (ACC), and 2 posterior cingulate cortex (PCC). Upon extracting the ICs, we estimated their dFNC by calculating Pearson’s correlation with a sliding tapered window. The window was created by convolving a rectangle with a 40-second step size with a Gaussian that had a standard deviation of 3. Interactions between individual ICs are represented as IC 1 / IC 2 (e.g., PCN 1 / ACC 1). For each participant, this resulted in data with 21 dFNC features and 124 time steps.

### C. Description of Clustering and Dynamical Feature Extraction

After extracting the dFNC for each study participant, we concatenated all dFNC samples and applied fuzzy c-means clustering. We clustered the data 50 times and selected the random seeds that had the highest fuzzy partitioning coefficient for their respective parameter combination. We used 5 clusters for easier comparison to previous studies that have uncovered 5 DMN dFNC clusters [2]. We optimized the fuzziness (i.e., m) parameter, considering values of 1.01 and 1.5. We also used an error of 0.0001 and maximum number of iterations of 1000. After assigning all samples to fuzzy states, we extracted dynamical features that have been used in previous studies to examine the effects of neurological and neuropsychiatric disorders upon brain activity [1], [5], [8]. Namely, we extracted the occupancy rate (OCR) for each study participant and state and the number of state transitions (NST) for each participant. Extracting these features required assigning each sample to the fuzzy state for which it had the highest probability of belonging.

### D. Description of Preexisting Feature Importance Approach

We applied an existing explainability approach for insight into the dFNC features most important to the clustering. We used Global Permutation Percent Change (G2PC) feature importance [4]. G2PC is an adaption of permutation feature importance to the domain of unsupervised clustering explainability and is broadly applicable to a variety of clustering algorithms. It has been applied for insight into dFNC and static FNC data in multiple studies [5], [6]. G2PC involves permuting (i.e., shuffling) a particular feature across samples for a number of repeats, reassigning the permuted samples to the previously identified clusters, and subsequently calculating the percent of samples that switch clusters following permutation. This process is repeated for each feature, and the features that result in the highest disruption in clustering are considered to be most important. We used 1000 repeats in our study.

### E. Description of Novel Explainable Clustering Approach

In this study, we developed a novel explainable clustering approach. While G2PC can be applied to fuzzy c-means, it does not take advantage of the probabilities of each sample belonging to each state. Our novel approach is highly similar to G2PC in that it involves permuting features and reassigning the permuted samples to the previously identified clusters. However, unlike G2PC it calculates the Kullback-Leibler divergence (KLD) between the probability distribution of each perturbed sample belonging to each cluster and the probability distribution of the corresponding unperturbed samples belonging to each cluster. The clustering is most sensitive to the features that result in the highest KLD in cluster probabilities when perturbed. We calculated the KLD for each sample following perturbation and then performed two subsequent calculations. We calculated the total KLD and the median KL divergence across samples for each feature for 1000 repeats. Additionally, to validate our assumption that our approach is able to capture the effects of perturbation upon more samples than G2PC, we perturbed each sample 100 times and calculated the percent of samples that had non-zero median KLD values for each feature.

### F. Statistical Analysis

We performed two sets of statistical analyses. (1) We wanted to quantify the similarity in feature importance estimates for G2PC and the total KLD and median KLD. To do this, we calculated the median importance for each feature based on each approach. We then ranked the features in order of median importance and performed pair-wise comparisons of the similarity of feature rankings for each approach with Kendall’s rank correlation. We obtained p-values for this analysis and applied FDR correction to reduce the likelihood of false positives. (2) We wanted to determine whether the states that we identified and their respective dynamical features were related to SZ. To that end, we performed two-tailed, independent sample t-tests comparing the OCRs and NSTs for SZs and HCs. We then performed false discovery rate (FDR) correction on the OCR p-values.

## III. RESULTS AND DISCUSSION

In this section, we describe and discuss our findings. We further discuss limitations and next steps for the study.

### A. Identification of 5 dFNC States

For easier comparison to previous studies [2], we identified 5 fuzzy states (i.e., clusters). Because we feature-wise z-scored the data before clustering, Figure 1 shows the re-scaled centroids of the fuzzy clusters. State 0 is characterized by low levels of positive connectivity. State 1 has high levels of positive connectivity in all domains except ACC/PCN, which has negative connectivity. State 2 has high levels of positive connectivity in all domains. State 3 is similar to state 1. However, it has slightly lower positive connectivity. Lastly, state 4 is similar to state 2 except for PCN3/ACC that has negative connectivity. Interestingly, the states that we identified seem to differ from those uncovered in [2], which is likely attributable to our use of z-scored dFNC features and fuzzy c-means.

**Figure 1.**
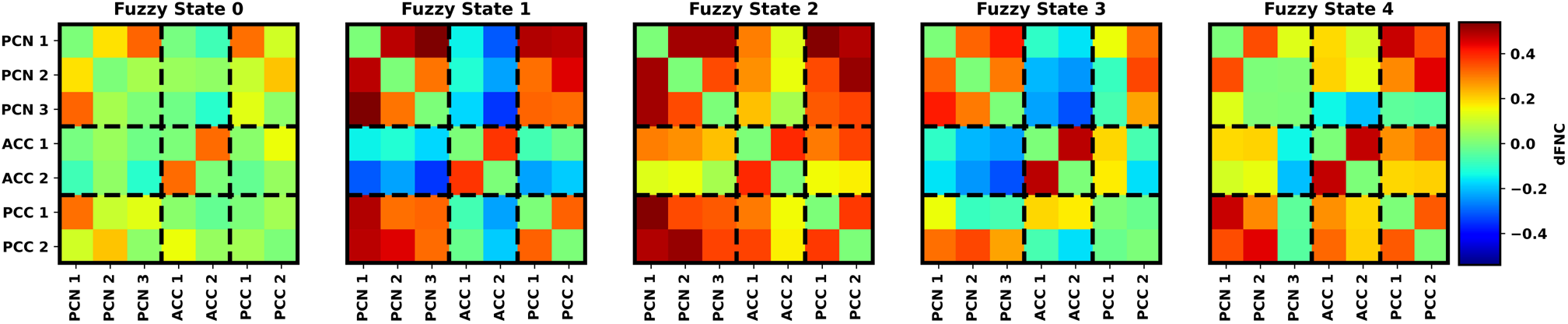
Fuzzy State Scaled Centroids. The scaled centroids for each state are arranged from left to right in ascending order. Each panel shares the same color bar to the right of the panel for Fuzzy State 4. The ICs associated with each dFNC feature are arranged on the x- and y-axes and are grouped based upon brain region (i.e., PCN, ACC, PCN).

### B. Identifying Top dFNC Features and Differences in Explanations

Figure 2 shows the explainability results for each of the approaches used in the study. G2PC, total GP2D, and median GP2D all identified PCN3/PCC1 and PCC1/ACC 2 as most important or among the most important features. PCN 3, PCC 1, and their interactions with other DMN nodes were also important to the clustering. At a high level, there was a high degree of overlap in importance across approaches. Based on Kendall’s rank correlation, there was a significant relationship between G2PC and total KLD (p < 0.05, τ = 0.35) without correction but not with correction. However, there was not a significant relationship between median GP2D and G2PC (τ = 0.17) and median GP2D and total GP2D (τ = 0.0). This finding was to be expected. Permutation typically only causes a minority of samples to completely switch clusters. As shown for G2PC, a maximum of around 10% of samples switched clusters completely. It also stands to reason that samples with a large enough change to switch clusters would also have a greater effect upon KL divergence that would be captured in a summation of KL divergence across samples (i.e., total GP2D). However, median KL divergence or GP2D, which was not correlated with either of the other approaches, is more likely to be representative of the permutation across samples, because as shown in the sample-level GP2D analysis of Figure 2, more than 70% of samples had non-zero KLD responses to perturbation.

**Figure 2.**
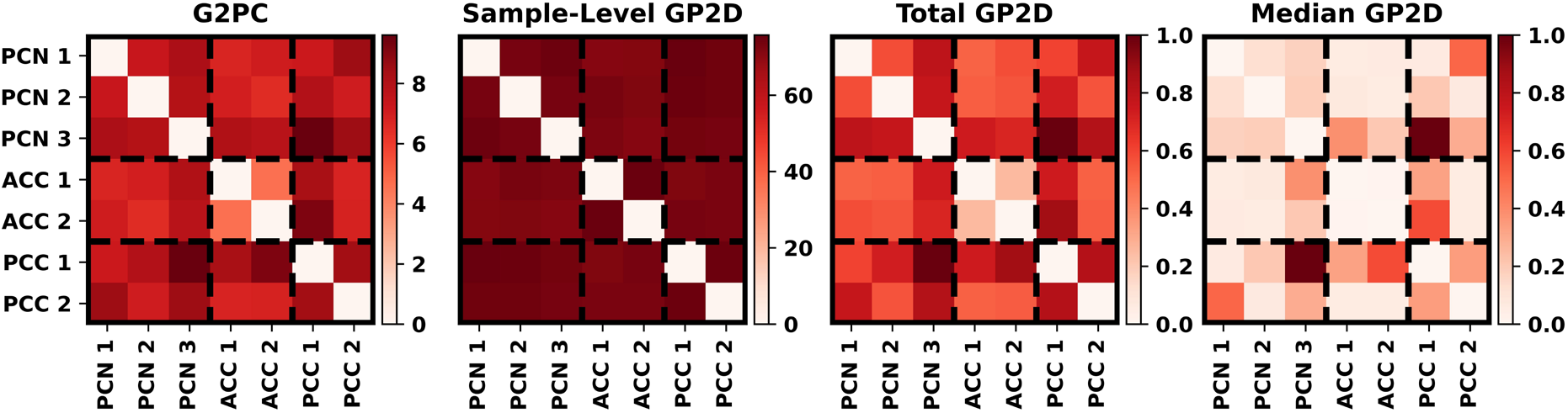
Explainability Results. From left to right, the panels show the G2PC, sample-level GP2D, total GP2D, and median GP2D results. The color bar corresponding to each panel is positioned to its right, and the ICs associated with each dFNC feature are arranged on the x- and y-axes and are grouped based upon brain region (i.e., PCN, ACC, PCN).

### C. Identifying Disease Related Differences in Extracted Dynamical Features

We extracted OCR and NST features related to the dynamics of the identified states. The features that we extracted are shown in Figure 3. There was not a statistically significant difference in NST between SZs and HCs. Additionally, with FDR correction, SZs spent significantly more time in state 1 than HCs (p < 0.01) and less time than HCs in state 3 (p < 0.001). Prior to correction, SZs spent significantly more time in state 2 than HCs (p < 0.05). Based on this, SZs spent more time in states of highly positive connectivity (i.e., states 1 and 2). In contrast, HCs spent more time in a state of moderate connectivity (i.e., state 3).

**Figure 3.**
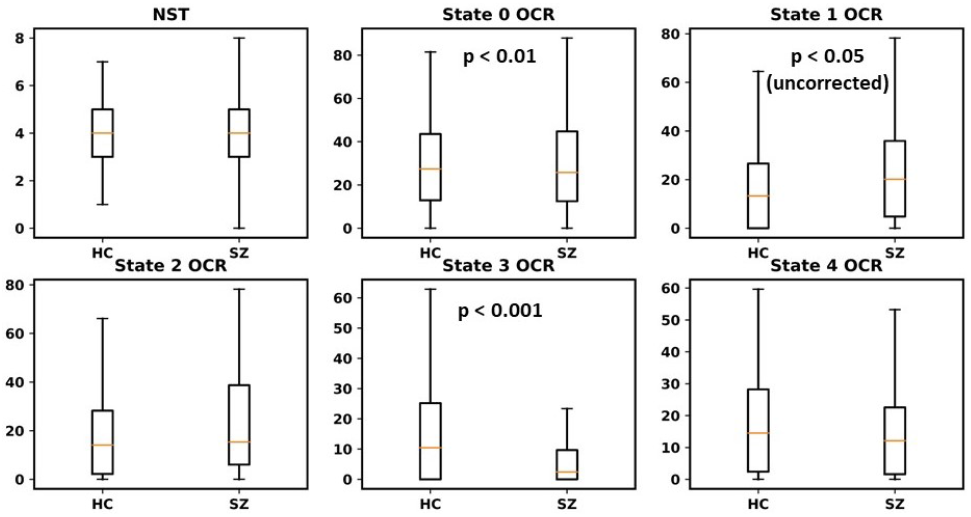
Comparison of Dynamical Features. Each panel shows a separate dynamical feature, with the top left panel showing NST and the other panels showing OCRs. HCs and SZs are on the left and right of each box plot, respectively. Features with significant differences between SZs and HCs have an accompanying p-value.

### D. Limitations and Next Steps

While we identified SZ-related differences in the dynamical features that we extracted, we confined our analysis to features that have been extracted in previous studies with hard clustering methods. In the future, we will present novel dynamical features that take advantage of the fuzzy clustering probabilities. Additionally, G2PC has a companion method called L2PC (i.e., Local Perturbation Percent Change) feature importance [4]. As we demonstrated in our sample-level GP2D analysis, it is possible to examine the effect of perturbation on a local basis with our approach, and future studies could explore the uses of a local permutation-based distribution divergence approach.

## IV. CONCLUSION

Existing explainability methods for identifying the importance of specific FNC features to identified clusters often use hard clustering and require that samples completely switch clusters. However, only a small portion of samples switch clusters, and resulting explanations are not representative of the overall clustering. In this study, we present GP2D, showing that it quantifies the effects of permutation upon the majority of samples and is thus able to more effectively quantify clustering feature importance than existing methods like G2PC. We demonstrate our approach within the context of DMN SZ analysis, uncovering differences in state dynamics between SZs and HCs and identifying specific intra-DMN interactions important to the clustering. We expect that our novel explainable clustering approach will enable further progress in rs-fMRI analysis and to other clustering applications.

## ACKNOWLEDGMENT

We thank those who collected the FBIRN dataset.

